# Deep Learning-Based Classification of CRISPR Loci Using Repeat Sequences

**DOI:** 10.1101/2024.06.27.601093

**Authors:** Xingyu Liao, Yanyan Li, Yingfu Wu, Xingyi Li, Xuequn Shang

## Abstract

With the widespread application of the CRISPR-Cas system in gene editing and related fields, the demand for detecting and classifying CRISPR-Cas systems in metagenomic data has continuously increased. The traditional classification of the CRISPR-Cas system mainly relies on identifying neighboring cas genes of repeats. However, in some cases where there is a lack of information about cas genes, such as in metagenomes and fragmented genome assemblies, traditional classification methods may become ineffective. Here, we introduce a deep learning-based method called CRISPRclassify-CNN-Att, which classifies CRISPR-Cas systems solely based on repeat sequences. CRISPRclassify-CNN-Att utilizes convolutional neural networks (CNNs) and self-attention mechanisms to extract features from repeat sequences. It employs a stacking strategy to handle sample imbalances across different subtypes and improves classification accuracy for subtypes with fewer samples through transfer learning. CRISPRclassify-CNN-Att demonstrates excellent performance in classifying multiple subtypes, particularly in subtypes with a larger number of samples. Although CRISPR loci classification primarily relies on cas genes, CRISPRclassify-CNN-Att offers a new approach as a significant complement to current methods. It can identify unclassified loci missed by traditional cas-based methods, breaking the limitations of traditional approaches, and simplifying the classification process. The proposed tool is freely accessible via https://github.com/Xingyu-Liao/CRISPRclassify-CNN-Att.

## Introduction

The CRISPR-Cas system, an adaptive immune defense mechanism, helps archaea and bacteria fend off viruses and plasmids, which consisting of leader sequences (often AT-rich), CRISPR array composed of repeats and spacers, and operons of CRISPR-associated genes responsible for encoding cas proteins^1, 2, 3, 4, 5, 6, 7^. The CRISPR-mediated response involves three main stages: spacer acquisition (adaptation), CRISPR RNA expression and maturation (processing), and interference with invading DNA (interference)^8, 9^. During the adaptation stage, the repeat serves as a template for integrating newly acquired spacer sequences to enhance immune defense, while in the expression stage, it facilitates the maturation of CRISPR RNA (crRNA), and in the interference stage, it plays a crucial role in binding crRNA to cas proteins^5, 10, 11, 12, 13, 14^. The ability of the CRISPR-Cas system to precisely recognize and cleave nucleotides makes it the most widely used tool in current genome engineering across biotechnology, bioprocessing, medicine, and agriculture, due to its low cost and efficiency.

CRISPR-Cas genes evolve rapidly in defense against invasion by mobile genetic elements, leading to their remarkable diversity^15^. Different subtypes of CRISPR-Cas systems exhibit significant differences in the composition of effector complexes, target nucleic acid types, and cleavage outcomes^16^. Currently, known CRISPR-Cas systems are classified into 2 classes, 6 types, and more than 40 subtypes based on the composition of effector complexes, the presence of signature, accessory cas proteins, and the architecture of CRISPR-Cas loci^17, 18^. Class 1 CRISPR-Cas systems include Types I, III, and IV, which are composed of multi-protein effector complexes. Class 2 consists of Types II, V, and VI, which are composed of single multi-domain effector proteins. Systems of Types I, II, and V identify and cleave DNA, while Type VI targets RNA, and Type III acts on both DNA and RNA for cleavage^18, 19^.

The classification of CRISPR-Cas systems relies on a gold standard of manual annotations by experts, published every few years^17, 20, 21^. The classification process typically involves identifying the CRISPR arrays, also known as repeats, and then searching the surrounding area for corresponding cas genes for classification, utilizing Hidden Markov Model searches or manual BLAST and PSI-BLAST searches^18, 22^. With the growing abundance of genomic sequences from uncultured microorganisms and metagenomic data, manual annotation has become challenging to maintain pace with. Currently, there are many attempts to design automated pipelines for identifying various elements of the CRISPR systems. For instance, CRISPRloci achieves comprehensive annotation of CRISPR-Cas systems, including the detection of CRISPR arrays, boundaries of cas cassettes, and classification of cas genes, based on machine learning methods^23^.

CRISPRcasIdentifier utilizes machine learning techniques to identify and categorize cas genes, enabling accurate annotation and classification of CRISPR-Cas systems^24^. CRISPRdisco determines the type of CRISPR systems by identifying CRISPR repeat sequences and cas proteins^25^. The classification methods for these CRISPR-Cas systems primarily rely on the accurate identification and categorization of cas genes. However, many CRISPR-Cas loci are complex, with approximately 40% exhibiting atypical structures, such as orphan arrays (lacking nearby cas genes) and distant arrays (located far from associated cas genes)^17, 26^. The absence of cas gene information can lead to the ineffectiveness of classification methods that rely on cas genes. Recent advances in sequencing technology accelerate the sampling and sequencing of metagenomes, generating vast amounts of publicly available data rich in microbial community information, serving as ideal sources for discovering novel CRISPR-Cas variants^27^. However, the complexity of these data not only increases the challenges for many assembly algorithms in handling repetitive sequences but also may result in the separation of CRISPR arrays from their corresponding cas genes in partially assembled loci^28, 29, 30, 31, 32^. The absence of cas gene information can lead to the ineffectiveness of classification methods that rely on cas genes. Due to the critical role of repeat sequences in the immune process of CRISPR-Cas systems, Russel et al. first proposed the CRISPRCasTyper, which is based on repeat sequences for the classification of CRISPR-Cas systems employing the XGBoost model^33^. On this foundation, Nethery et al. validate and expand this technique by exploring new model input features and analyzing the contributions of underlying features for classification of CRISPR-Cas systems^34^. Currently, deep learning methods excel in several fields, automatically extracting features from sequence data, capturing subtle patterns and deep information, thereby significantly improving task accuracy and effectiveness compared to traditional machine learning methods.

Here, we propose a novel deep learning-based approach, CRISPRclassify-CNN-Att, that achieves classification of CRISPR-Cas systems based on repeats. We use convolutional neural networks (CNNs) to capture local patterns and self-attention mechanisms to focus on essential features when processing the input data, including encoded repeat sequences and extracted features such as *k-mer* frequencies, GC content, and sequence lengths^34^. Due to the imbalance in sample sizes across subtypes in the dataset, we adopt a stacking strategy^35^. We train separate models for subtypes with more and fewer samples as base model, utilizing transfer learning to improve classification accuracy for rare subtypes, and then use the outputs of these models combined with extracted features as inputs for the meta-classifier, XGBoost^36, 37^. This stacking strategy improves feature extraction efficiency and accuracy, achieving precise classification of CRISPR-Cas systems. Additionally, based on the base model, we analyze the relationship between *k-mers* and subtypes.

Through ablation experiments, we validate the positive impact of sequence features, including *k-mer* frequency, GC content, and sequence length, on the model’s performance.

CRISPRclassify-CNN-Att is the first to utilize deep learning for CRISPR-Cas system classification based on repeat sequences, thereby enhancing classification accuracy and expanding subtype coverage. The flowchart is shown in Figure 1. Direct use of repeat sequences simplifies the classification process compared to traditional methods relying on cas genes. Moreover, even in the absence of adjacent cas gene information, classification can still be achieved through repeat sequences, thus overcoming limitations of traditional methods in identifying CRISPR-Cas systems using cas genes.

**Figure 1.**
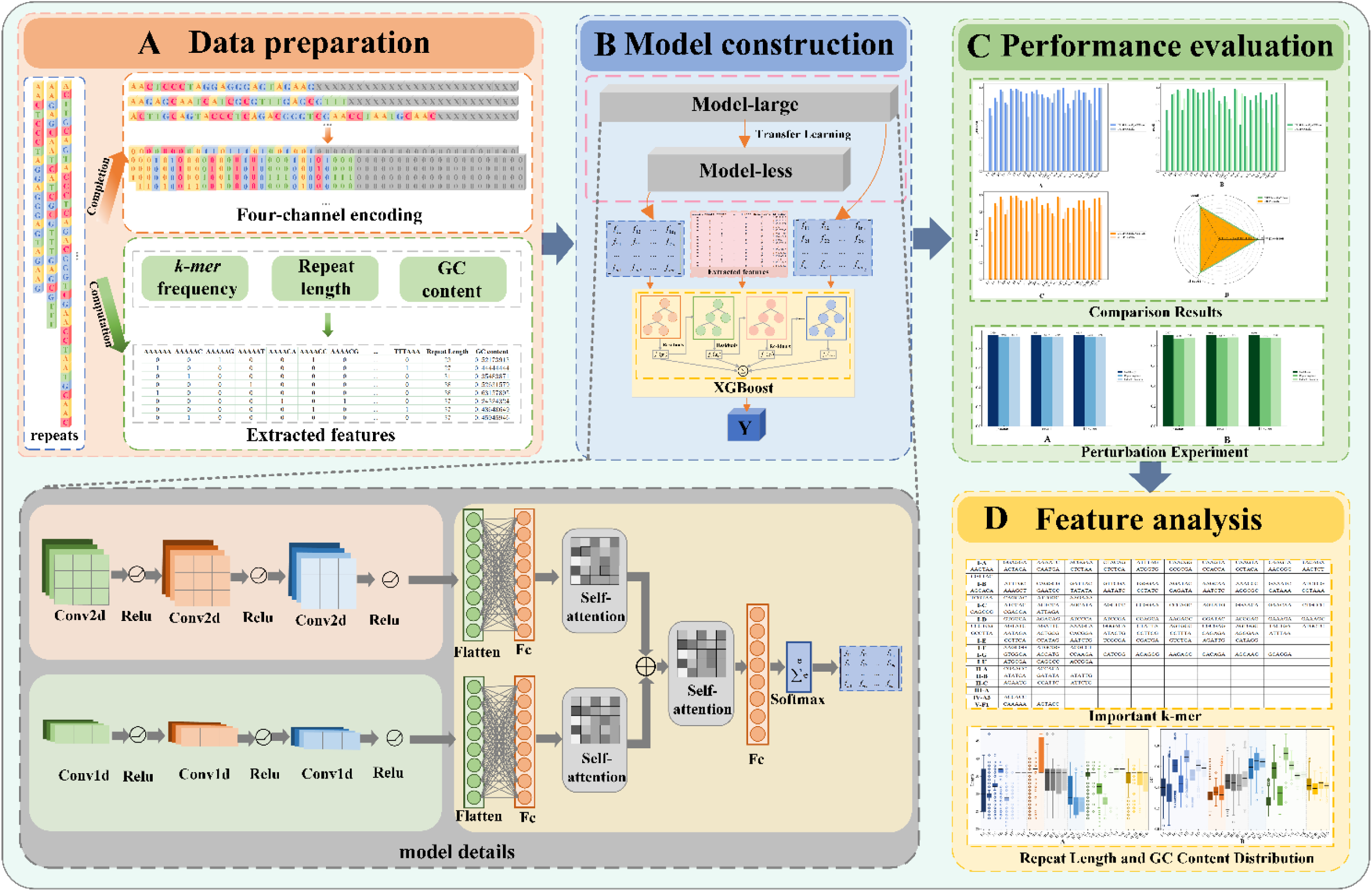
The workflow of CRISPRclassify-CNN-Att. (A) The data consists of two parts: one part is the encoded repeat sequences, and the other part includes features extracted from the repeats (*k-mer* frequency, repeat length, GC content). (B) Model-large (for subtypes with a large number of samples) and Model-less (for subtypes with less sample data) have their detailed structures shown in the gray section. (C) The performance evaluation includes comparative experiments and ablation experiments. (D) The feature analysis includes identifying important *k-mers* and examining the distribution of repeat length and GC content.

## Results

### CRISPRclassify-CNN-Att achieves accurate classification of CRISPR-Cas systems

CRISPRclassify-CNN-Att employs deep learning techniques, integrating repeat sequences and extracted features, including *k-mer* frequencies, repeat length, and GC content, to achieve the classification of CRISPR-Cas systems. The *k-mer* frequency provide detailed information about sequence composition, sequence length offers important information about the scale of the sequences, and GC content reflects the nucleotide composition characteristics. By combining these multidimensional data, CRISPRclassify-CNN-Att achieves accurate classification of CRISPR-Cas system. We train separate models for subtypes with large and less samples (Model-large and Model-less), integrate these models using a stacking approach, and employ transfer learning to fine-tune the model for subtypes with less samples based on the model for subtypes with large samples, thereby improving classification capability. Model-large (the model for subtypes with large samples) comprises 11 subtypes (I-A, I-B, I-C, I-D, I-E, I-F, I-G, II-A, II-C, III-A, V-A), while Model-less (the model for subtypes with less samples) encompasses 11 subtypes (I-U, II-B, IV-A3, V-B1, V-F1, V-F2, V-K, VI-A, VI-B1, VI-B2, VI-D), totaling 22 subtypes.

To evaluate CRISPRclassify-CNN-Att, we compare it with CRISPRclassify, an XGBoost-based repeat classification method for CRISPR-Cas systems. For base models (Model-large and Model-less), we use half of the dataset as the training set and the remaining half as the test set. The outputs of the base models on the test set form a new dataset, which serves as both the training and test sets for the meta-model, with a 1:1 split for training and testing. The evaluation metrics include precision, recall, and f1-score for both the overall performance and each individual subtype. Figure 2 shows the detailed evaluation results and Table S1 presents the specific data. We find that CRISPRclassify-CNN-Att demonstrates higher precision, recall, and f1-score in most subtypes compared to CRISPRclassify, especially in those with larger sample sizes. Specifically, for precision, CRISPRclassify-CNN-Att outperforms CRISPRclassify in 90.9% of the subtypes; for recall, CRISPRclassify-CNN-Att exceeds CRISPRclassify in 86.4% of the subtypes; and for f1-score, CRISPRclassify-CNN-Att surpasses CRISPRclassify in 90.9% of the subtypes. These results indicate that CRISPRclassify-CNN-Att exhibits superior performance in the classification of CRISPR-Cas systems based on repeat sequences, providing a more reliable method for CRISPR-Cas system classification.

**Figure 2.**
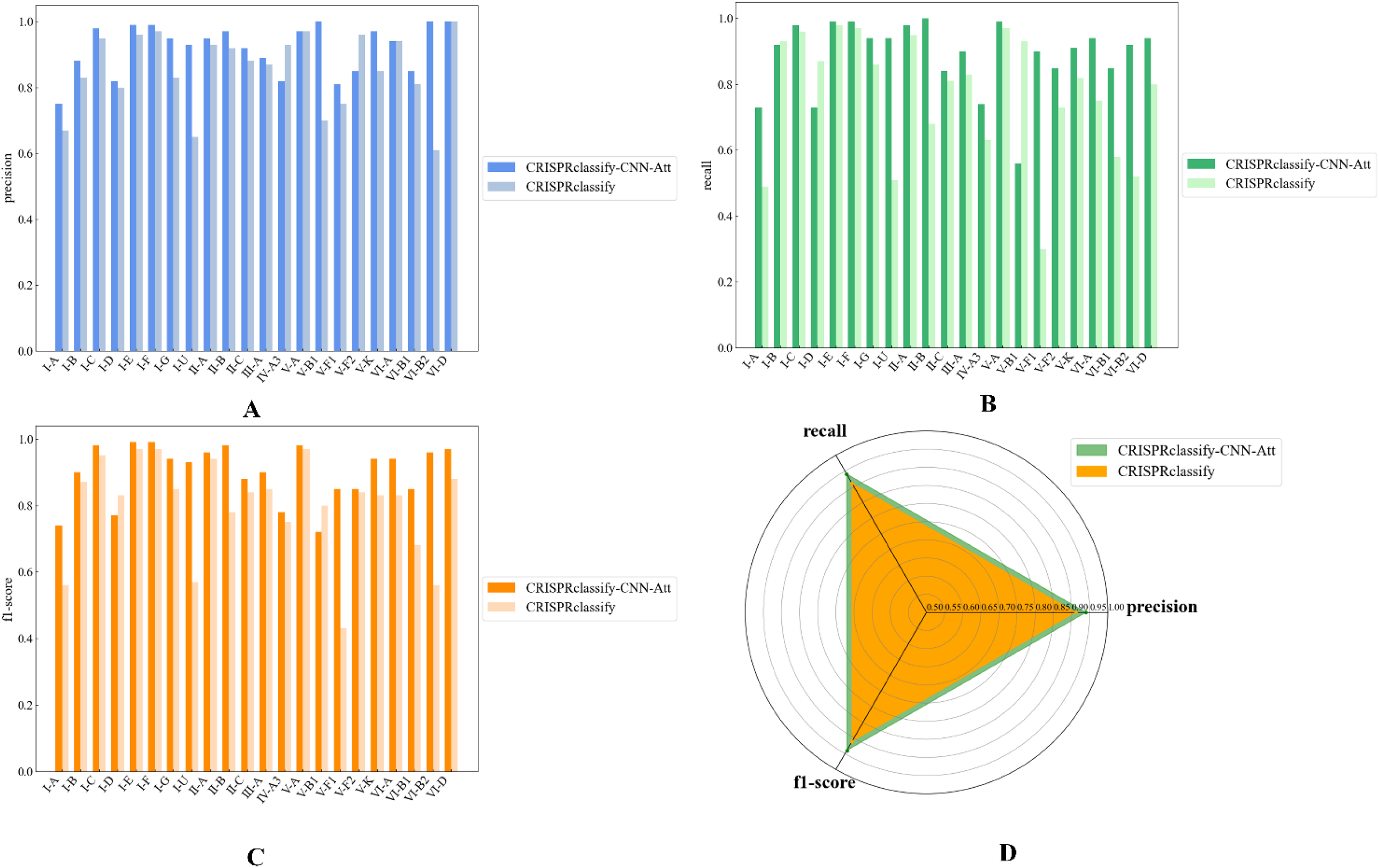
Performance evaluation is divided into individual subtype and overall assessment. (A), (B), and (C) are the comparative results for precision, recall, and f1-score on individual subtypes. (D) represents the overall metric comparison.

During the model training process, we observe that the sample sizes for subtypes III-B and III-C are relatively large, yet the model consistently performs poorly in classifying them. We exclude the influence of sample quantity and explore potential reasons. We calculate the similarity between repeat sequences of subtypes with more samples, specifically the normalized minimum Levenshtein distance between repeat sequences of subtype *s_i_* and subtype *s_j_*, and also compute the Euclidean distance between their additional features, as depicted in Figure 3. We find that the repeat sequence similarity between subtypes III-B/C and certain subtypes within type I systems is significantly higher than the similarity between other subtypes, and the Euclidean distances between the extracted features are also smaller than those of other subtypes. Existing research indicates that III-B/C systems typically lack independent adaptation mechanisms and instead rely on selecting the necessary cas proteins from other genomic loci encoding CRISPR-Cas systems to expand their CRISPR arrays^38^. When a strain contains both III-B or III-C loci lacking cas1/2 genes and a type I system, the repeat sequences of the III locus are highly similar to those of the type I locus. This is because the type I cas1-cas2 adaptation complex recognizes and processes the III repeat sequences^20, 39, 40^. In contrast, when III-B/C systems contain cas1/2 genes, the similarity between the repeat sequences of the III and type I loci significantly decreases, indicating that III systems have their own independent cas1-cas2 complex. In summary, the repeat sequences of III-B/C subtypes lacking cas1/2 genes are easily misclassified as type I due to their high sequence similarity to type I systems.

**Figure 3.**
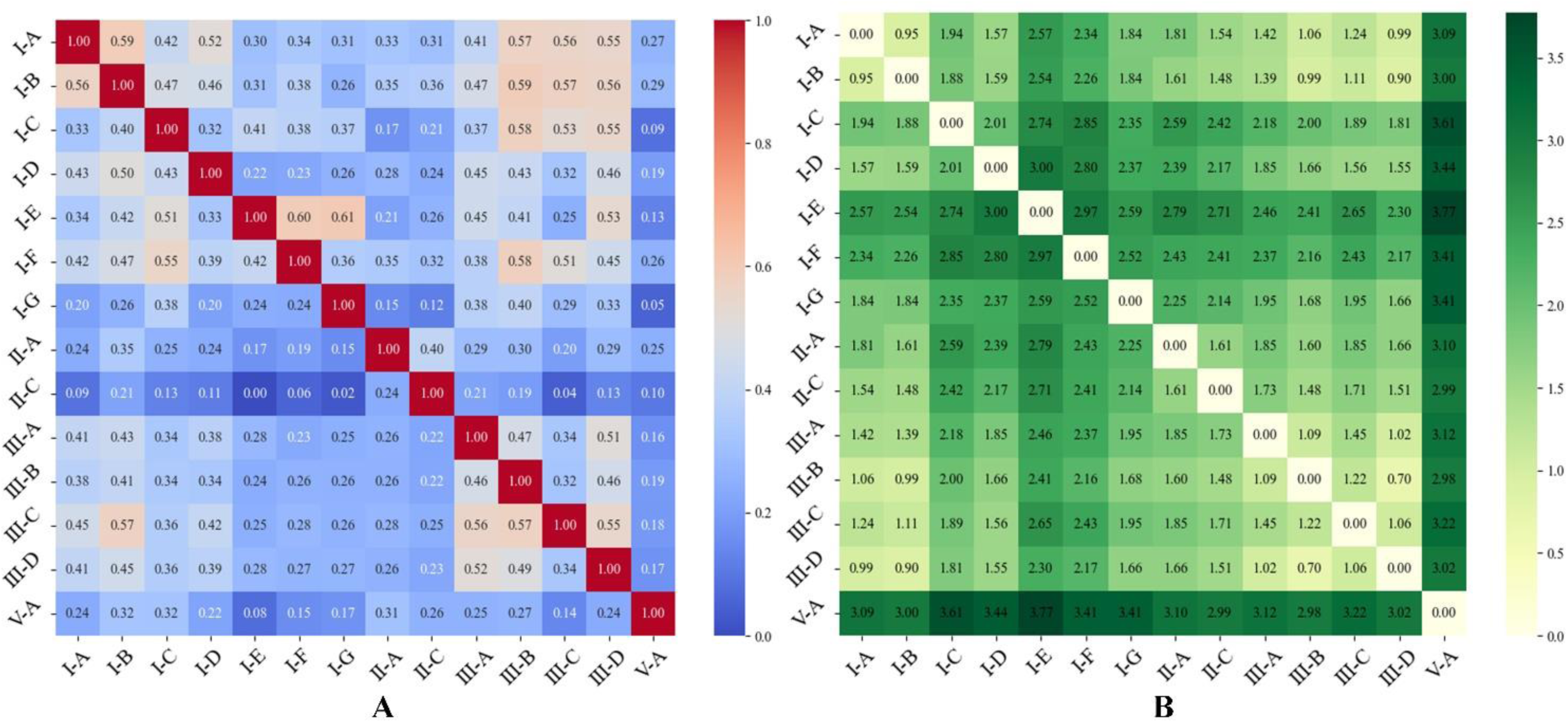
(A) shows the sequence similarity between subtypes with a large number of samples, defined as the normalized minimum Levenshtein distance between subtype sequences. (B) shows the Euclidean distance between extracted feature vectors of subtypes with a large number of samples.

### The impact of repeat sequence and extracted features on model performance

CRISPRclassify-CNN-Att takes two components as input: one is the encoded repeat sequence, and the other consists of extracted features, including *k-mer* frequency, sequence length, and GC content. To explore the impact of the repeat sequence and extracted features on model performance, we conduct ablation experiments based on the base model. During the experiments, we individually remove the sequence and extracted features, use each feature as input separately, and observe the effects of feature ablation on model accuracy, recall, and f1-score. Figure 4 respectively displays the comparison between Model-large and models utilizing only encoded repeats or extracted features, as well as between Model-less and models utilizing only encoded repeats or extracted features. Table S2 presents the specific data. The experimental results demonstrate that repeat sequence significantly affect model performance, underscoring the critical role of sequence information in CRISPR-Cas system classification. Additionally, we find that the extracted features, including sequence length, *k-mer* frequency, and GC content, are equally crucial for enhancing classification efficacy. These findings further highlight the essential role of multidimensional data integration in this classification system, providing valuable insights for model optimization and improvement.

**Figure 4.**
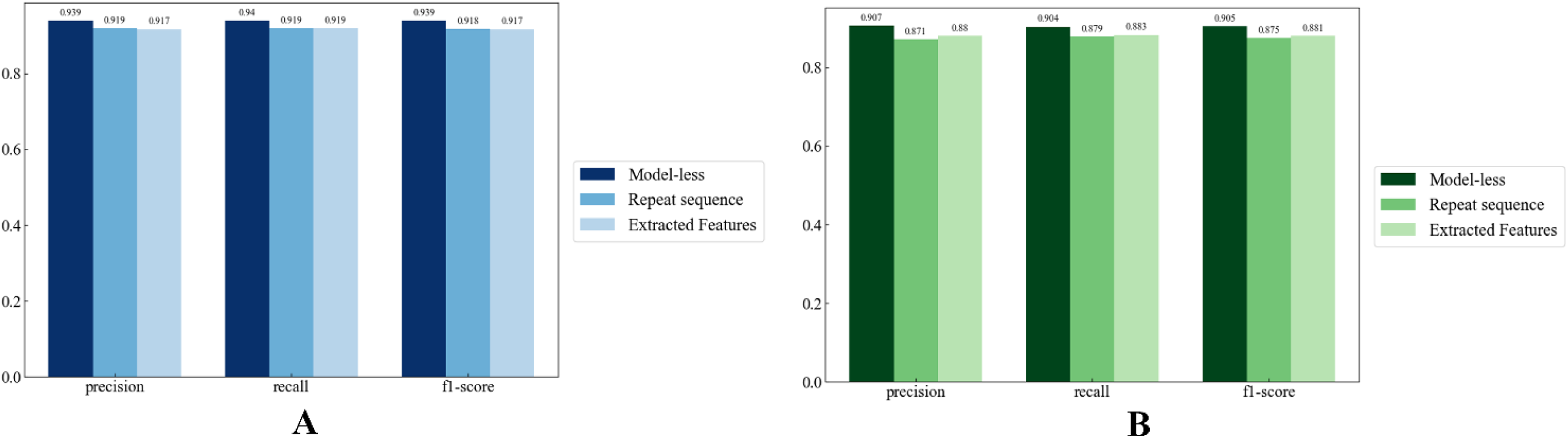
The comparative results of feature ablation studies for Model-large (A) and Model-less (B). It illustrates the effects of removing individual repeat sequences and extracted features on model performance, including accuracy, recall, and f1-score.

### Important *k-mer* in Each CRISPR-Cas System subtypes

The frequency of *k-mer* occurrences is a set of key features containing important information about the sequence. We use the base model to analyze the important *k-mers* in each CRISPR-Cas system subtype. To identify these key *k-mers*, we systematically set the occurrence frequency of each *k-mer* in the test set to zero and observe changes in the model’s prediction metrics for each subtype, including precision, recall, and f1-score. If setting a particular *k-mer* to zero results in a decrease in prediction performance, that *k-mer* is considered important for that subtype. Using this method, we identify the *k-mers* that significantly contribute to the classification of each subtype, reflecting the sequence specificity of different subtypes. Table 1 shows the important *k-mers* identified in each subtype. Table 2 further illustrates the unique key *k-mers* for each subtype, helping us better understand the sequence specificity differences among the subtypes. For example, "GGAGGA" "AAAATC" and "ACGGAA" uniquely contribute to the prediction of subtype I-A, while "ATTTGC" "CAGGCG" and others are unique contributors to subtype I-B. We calculate the average occurrence frequency of *k-mers* within each subtype and find a high overlap with the key *k-mers* identified by the model based on *k-mer* occurrence frequency. This suggests that frequently occurring *k-mers* play a crucial role in the classification task. These findings further validate the accuracy of the model’s selection of key *k-mers* and underscore their significance in CRISPR-Cas system classification. These key *k-mers* contribute positively to CRISPR-Cas system classification, enhancing model performance, interpretability, and aiding in the identification of sequence specificity among different subtypes.

**Table. 1.**
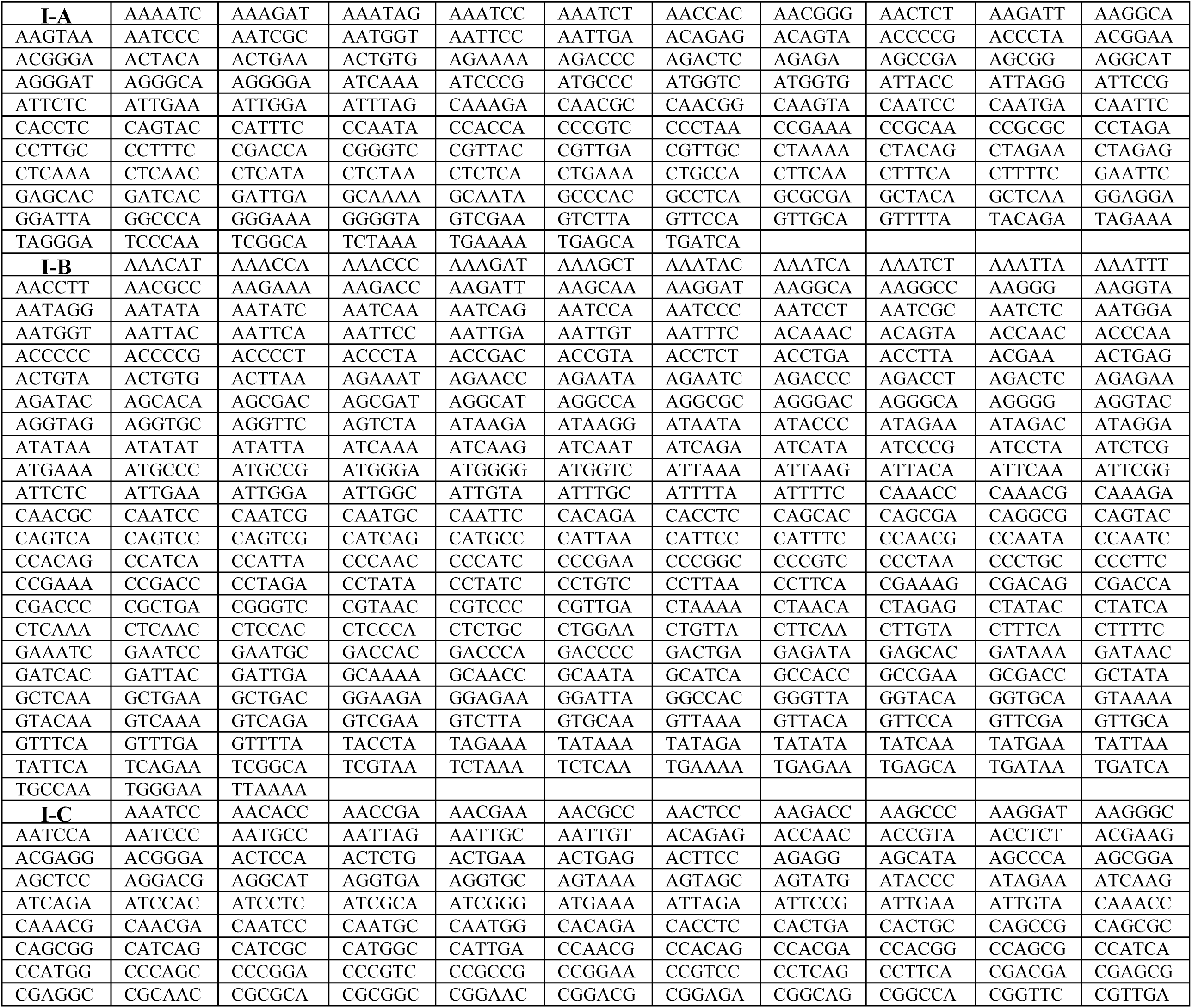
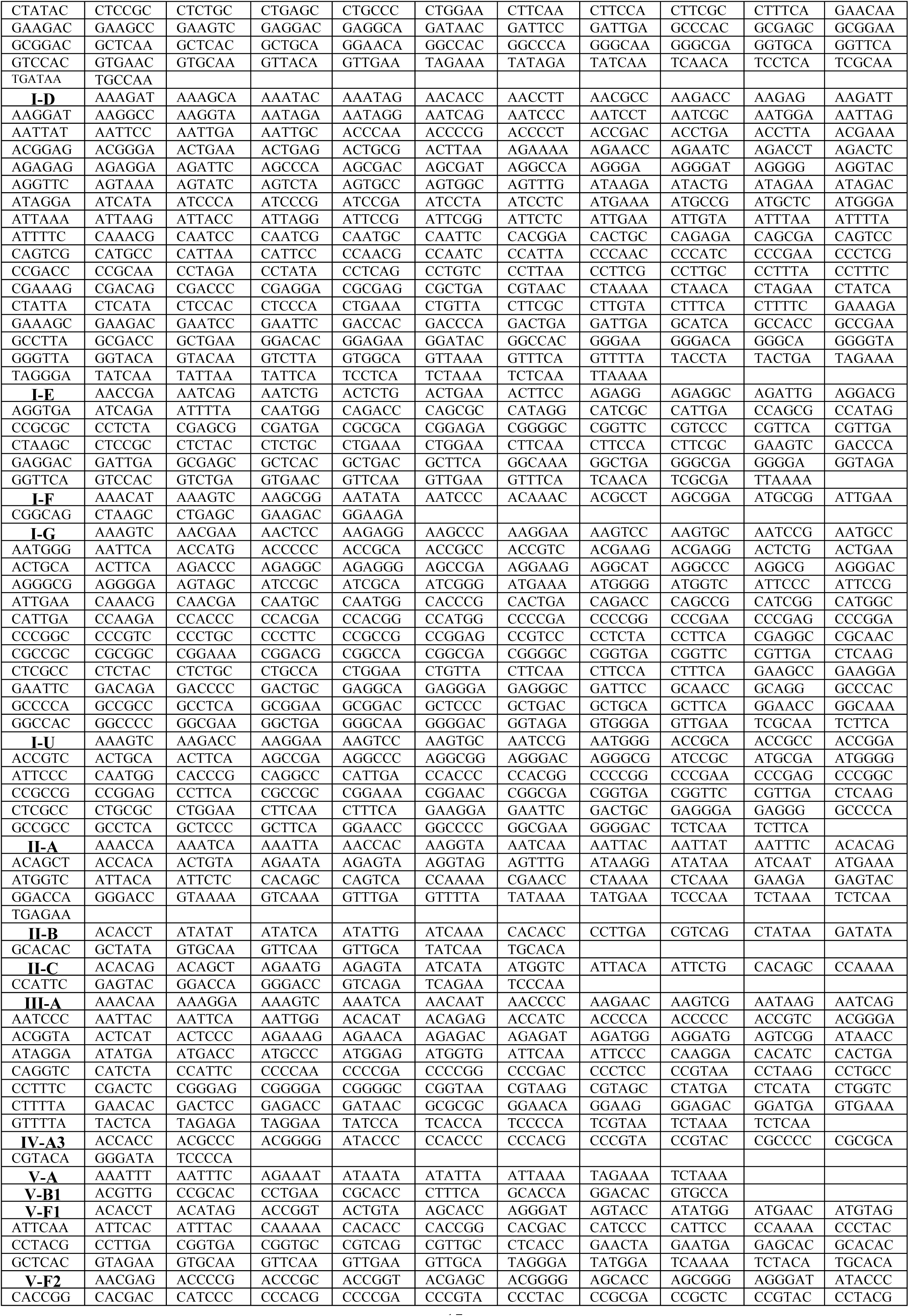
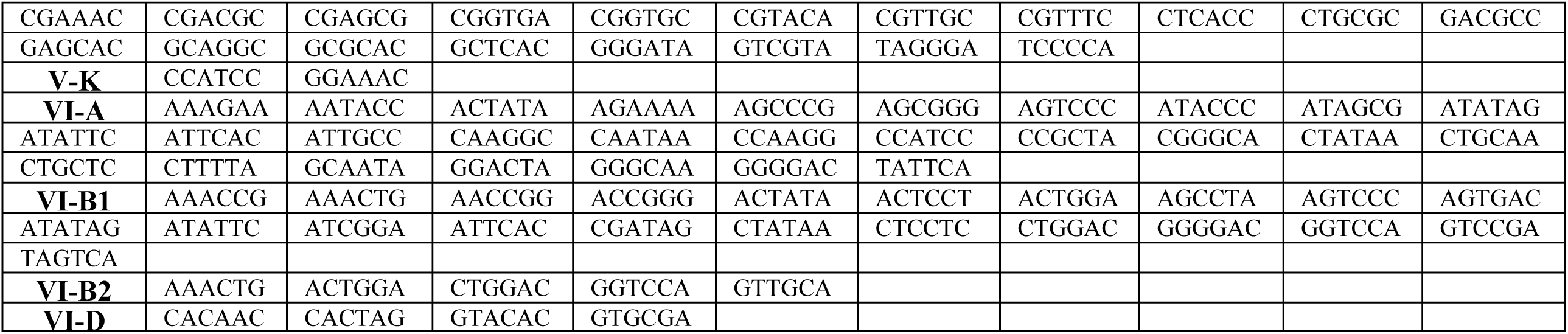
All the important *k-mers* within each subtype, setting the *k-mer* frequency to zero negatively impacts the predictive performance.

**Table 2.**
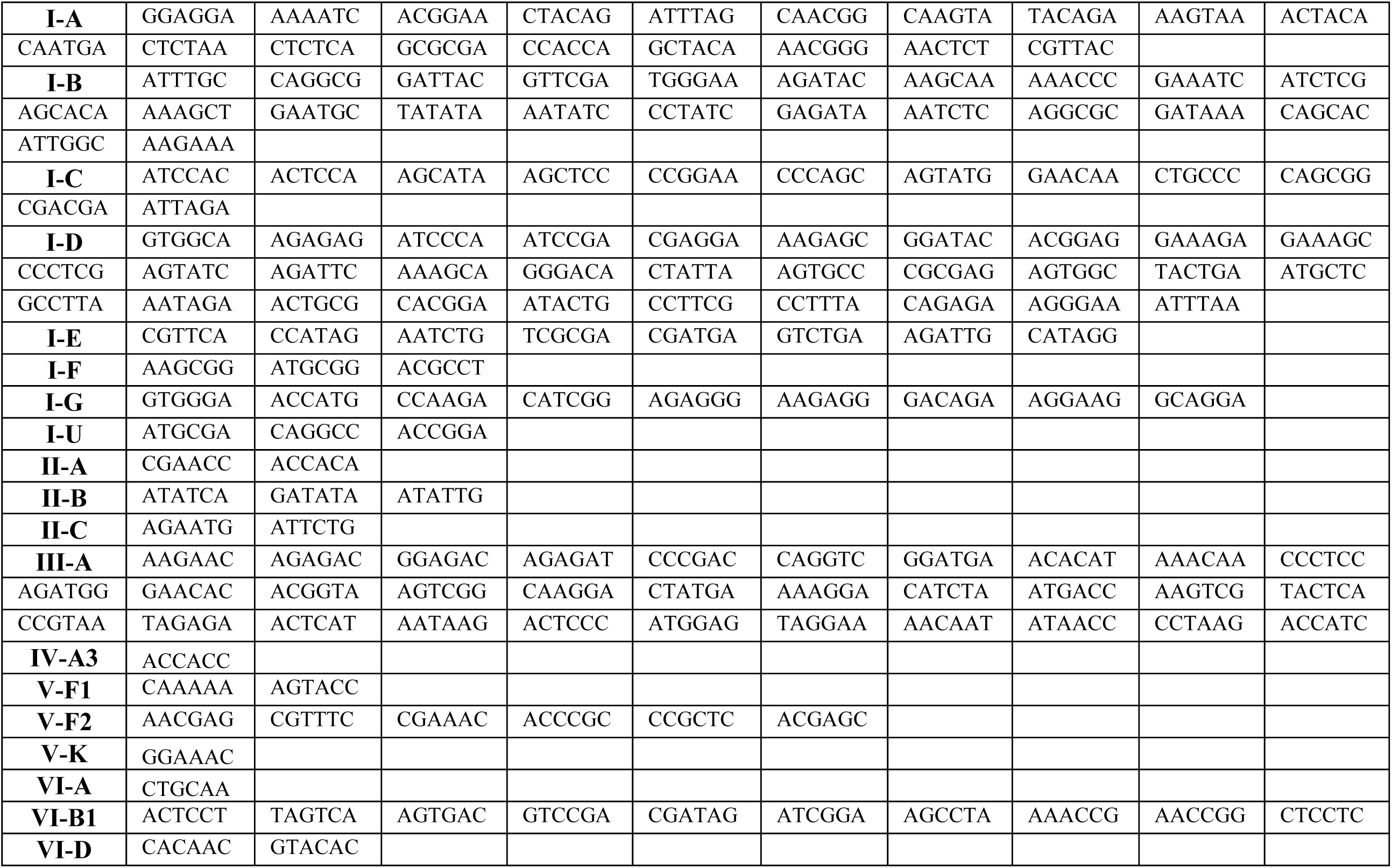
Unique important k-mers exist exclusively within individual subtypes.

Furthermore, we identify the important *k-mers* in each subtype from the perspective of the repeat sequence. We replace the encoding of each *k-mer* appearing in the sequence with [0,0,0,0] one by one, then observe changes in the model’s performance metrics, including precision, recall, and f1-score. If replacing the encoding of a specific *k-mer* with [0,0,0,0] results in a significant drop in prediction performance, that *k-mer* is considered important for that subtype. Table S2 lists the important k-mers identified from the repeat sequence perspective and Table S3 shows the unique important k-mers for each subtype. We compare the important *k-mers* obtained from the *k-mer* occurrence frequency for each subtype with those obtained from the sequence perspective and identify some common *k-mers*. Table 3 showcases these common *k-mers*. These *k-mers* reflect the sequence specificity of different subtypes and play a crucial role in the classification task. This not only helps improve the model’s performance and interpretability but also provides valuable insights into the sequence characteristics of each subtype. Additionally, these common *k-mers* offer a valuable reference for future research, aiding in the further exploration of the functions and mechanisms of the CRISPR-Cas system.

**Table 3.**
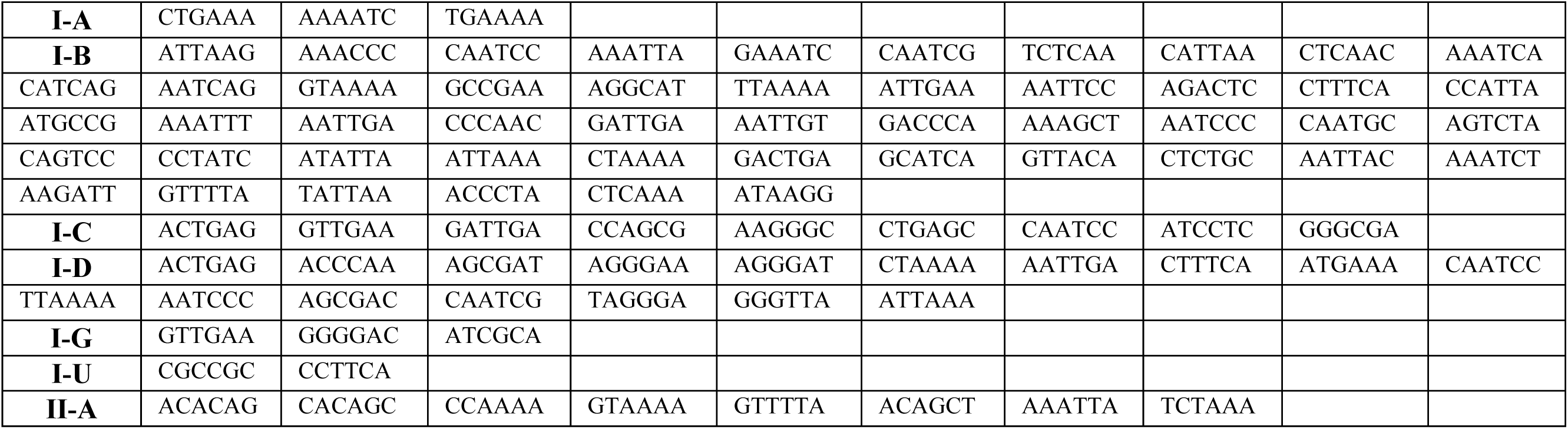
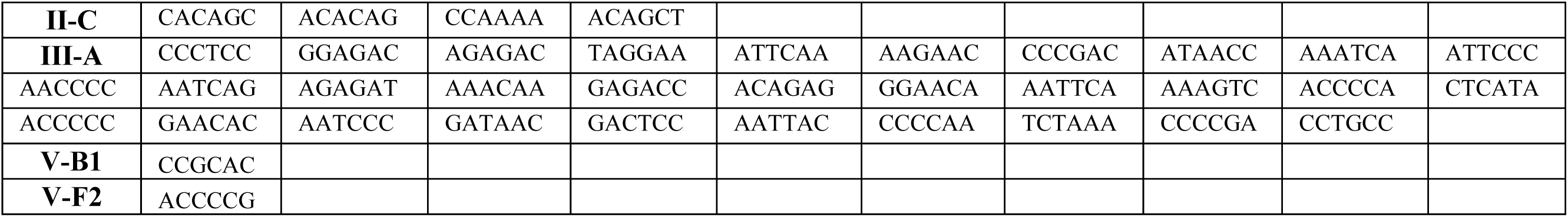
The common important *k-mers* for each subtype based on *k-mer* frequency and repeat sequence perspective.

In addition to *k-mer* frequency, sequence length and GC content also play a significant role in model performance. We calculate the distribution of these features for each subtype, as shown in Figure 5. In type I CRISPR-Cas systems, the repeat sequence lengths vary significantly among the I-A to I-E subtypes, while I-F and I-G have conserved repeat lengths of 28 bp and 36 bp. In type II systems, the lengths of repeat sequences are relatively conserved. For instance, subtype II-A has a conserved repeat length of 36 bp, while subtype II-B has a conserved repeat length of 37 bp. Although the distribution of repeat lengths in type III systems is relatively dispersed, their interquartile range distributions are similar, with a median of approximately 35 or 36 bp. The repeat lengths in type IV systems vary greatly, with only the interquartile range distributions of IV-A1, IV-A2, and IV-A3 being similar. The repeat lengths in type V systems are widely distributed, with median lengths ranging from 25 bp (V-D) to 37 bp (V-E). The quartile range distribution in type VI systems is similar, with a median length of approximately 36 or 37 bp. The GC content distribution varies significantly among different subtypes. Type I (I-A to I-U) exhibits a wide range of GC content, with some subtypes like I-A and I-B having lower median GC content, while others like I-G have higher GC content. Type II (II-A to II-C) has a more concentrated median GC content, with II-A and II-B being higher, and II-C relatively lower. Subtype III (III-A to III-D) has relatively lower GC content, especially III-C and III-D. Type IV (IV-A1 to IV-A3) shows higher median GC content and wider distribution. Type V (V-A to V-K) displays a wide range of GC content, with V-A and V-B1 having higher GC content, while V-G and V-J have lower content. Type VI (VI-A to VI-D) has a relatively lower GC content distribution, with VI-B1 and VI-B2 having lower median GC content.

**Figure 5.**
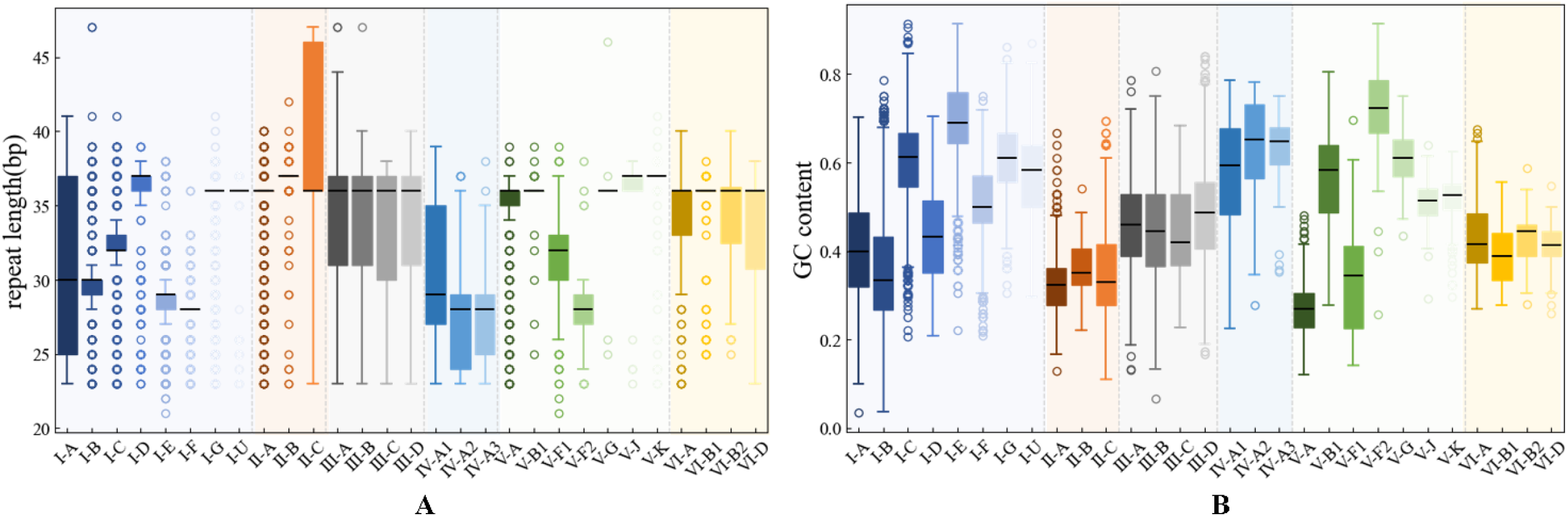
(A) showcases the distribution of repeat length for subtypes with a sample size greater than 30, while (B) displays the distribution of GC content.

## Discussion

In this work, we propose a novel deep learning-based method for classifying CRISPR-Cas systems based on repeat sequences. We use convolutional neural networks (CNNs) to process repeat sequences, automatically extracting and learning important features. We incorporate self-attention mechanisms to emphasize key features, enhancing the model’s understanding and classification capability. To improve classification performance, we integrate extracted features such as *k-mer* frequency, GC content, and sequence length, providing more comprehensive information for distinguishing different types of CRISPR-Cas systems. Due to the significant variation in sample sizes among different subtypes, the model tends to favor classes with more samples. Therefore, we train separate models for subtypes with large and less samples. When training models for subtypes with fewer samples, the model struggles to learn adequately due to the limited sample size. To address this, we employ transfer learning, first training a base model on subtypes with large samples and then fine-tuning it on subtypes with fewer samples to enhance performance. Our method relies solely on repeat sequences, thereby simplifying the classification process and overcoming the limitations of traditional methods dependent on cas genes. This approach provides a more efficient and accurate method for the classification of orphan and distant arrays.

Accurately classifying III-B/C subtype CRISPR-Cas systems presents challenges based on repeat sequences. During the training of models, when datasets include both III-B/C subtypes and type I systems, the evaluation metrics for both III-B/C and type I systems significantly decrease, with the precision of III-B/C systems dropping below 50%. Excluding sample size as a factor, we investigate potential reasons. Research suggests that III-B/C systems, lacking independent adaptation mechanisms, expand their CRISPR arrays by selectively acquiring necessary cas proteins from CRISPR-CAS systems encoded at other genomic loci^38^. When the cas1 and cas2 genes are absent from the locus, the repeat sequences of III loci bear high similarity to those of type I loci, leading to confusion and a high probability of misclassification of III-B/C as commensal type I systems^34^. However, when III-B/C loci contain cas1/2 genes, the similarity between the repeat sequences of III and type I loci significantly decreases, indicating the presence of an independent cas1-cas2 complex in III-B/C loci. CRISPR arrays and cas genes co-evolve as two distinct modules within the CRISPR-Cas system, with intricate interconnections and recombination events facilitating adaptive immunity. The tight associations and recombination between adaptive modules add complexity to the classification of CRISPR-Cas loci^18^.

The repeat sequences and extracted features (such as *k-mer* frequency, GC content, and sequence length) play a crucial role in the model’s performance. A systematic exploration is conducted on the base model through ablation experiments, revealing the pivotal role of sequence features in capturing subtle differences among CRISPR-Cas system subtypes. These sequence features assist the model in accurately distinguishing between different types of CRISPR-Cas systems, thereby enhancing classification capability. Furthermore, the incorporation of additional features further improves the model’s performance. By incorporating extra information such as *k-mer* frequency, GC content, and sequence length, the model gains a more comprehensive understanding of sequence characteristics and can better classify different subtypes of CRISPR-Cas systems. This integrated use of sequence and ectracted features enriches the model’s feature space, thereby enhancing its ability to classify CRISPR-Cas systems and improving the reliability and robustness of classification results.

In different subtypes of CRISPR-Cas systems, the distribution of repeat sequence lengths exhibits distinct characteristics. Within each subtype, the repeat lengths are conservative^25^. For example, in type I systems, the repeat lengths of I-F and I-G remain at 28 bp and 36 bp, respectively. In type II systems, the repeat length of subtype II-A is 36 bp, while subtype II-B maintains a length of 37 bp. Although the distribution of repeat lengths in type III systems is relatively dispersed, their quartile distributions are similar, with a median of approximately 35 or 36 bp. The quartile distributions in type VI systems are also similar, with a median length of approximately 36 or 37 bp.

## Methods

### The overall framework of CRISPRclassify-CNN-Att

CRISPRclassify-CNN-Att is a deep learning-based method for classifying CRISPR-Cas systems based on repeats, comprising the following steps: data preparation, model construction, performance evaluation, and feature analysis.

Data preparation consists of two parts. The first part involves processing repeats by padding and performing one-hot encoding to handle uneven lengths. The second part involves extracting additional features from repeats, including a *k-mer*-based feature set (the occurrence frequencies of each distinct *k-mer* in the repeat sequence and its reverse complement), as well as a set of biological features (repeat length, GC content).

We construct a classification model based on CNN^41^ and self-attention mechanism^42^. To address the imbalance in sample sizes across subtypes in the dataset, we adopt a stacking strategy^35^. Separate models are trained for subtypes with large and less sample sizes, and their outputs and extracted features are incorporated as new features into the meta-classifier, XGBoost^36^. This approach effectively leverages the predictive capabilities of each model and synergizes their strengths to improve overall classification performance.

We evaluate the model performance using precision, recall, and f1-score, which includes both overall performance and performance for each individual class. Additionally, based on the base model, we analyze the relationship between *k-mer* and subtypes to identify crucial *k-mers* within each subtype.

### Data preparation

The data sources are divided into two parts: one part from Russel et al.^33^, which includes 20,142 repeat sequences involving 44 subtypes, and the other from Nethery et al.^34^, which includes 15,646 repeat sequences involving 30 subtypes. The initial sources of these two datasets are the selected datasets of Makarova et al. and Pinilla-Redondo et al. ^18, 43^, which contain genomes with classified CRISPR loci^23^, downloaded from the National Center for Biotechnology Information (https://www.ncbi.nlm.nih.gov/). The repeat sequences were extracted using the MinCED tool (github.com/ctSkennerton/minced) a tool derived from CRT with default options^44^. After deduplication and filtering for repeats with sample sizes greater than 30, the dataset contains 25 subtypes and 28,388 repeat sequences.

The input data consists of two parts. The first part involves processing the repeat sequences, including padding, and encoding. Since the repeat sequences vary in length from 21 to 48 bp^27, 45^, we use ’X’ to uniformly pad the sequences to 48 bp. Then, we apply one-hot encoding to the padded sequences, with ’X’ represented as [0,0,0,0], resulting in four-channel sequence encoding. The second part involves extracting an additional set of features from the repeat sequences, including a *k-mer*-based feature set composed of the occurrence frequencies of each distinct *k-mer* in the repeat sequence and its reverse complement, as well as a set of biological features including the repeat length, the frequencies of G and C nucleotides in the repeat sequence, and the nucleotide matching score between the repeat sequence and its reverse complement at the same index positions.

### Model construction

We construct a robust multi-classification model leveraging Convolutional Neural Networks (CNN) ^41^augmented with a self-attention mechanism^42^ to enhance feature representation and capture important patterns in the data. The input data for the model consists of two parts: encoded sequences and additional extracted features(repeat length and GC content), which are processed through separate convolutions. This approach ensures a comprehensive analysis of both sequence data and biological features, thereby significantly enhancing classification performance.

For the four-channel encoded sequence data, we apply a two-layer convolution with ReLU activation, while the extracted features undergo three layers of convolution with ReLU activation.

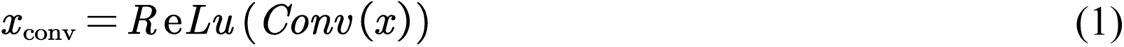

The definition of the ReLU function is:

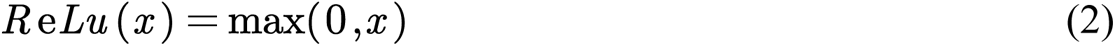

After passing through multiple convolutional layers, the features are flattened, and then feature weights are adjusted using a self-attention mechanism to focus on important information:

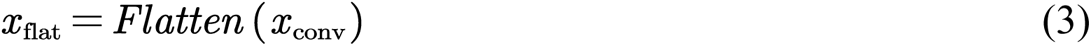

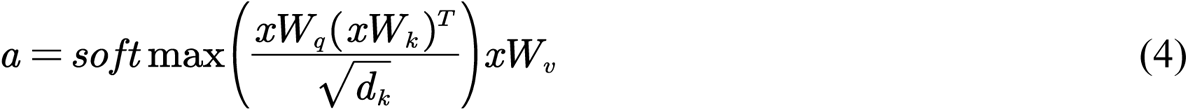

where *W_q_*, *W_k_* and *W_v_* represents three weight matrices used to map the input *x* to query, key, and value spaces. 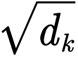 is a scaling factor used to prevent the dot product result from becoming too large. After self-attention processing, the features merge and enter the final fully connected layer, where they are transformed into probabilities using the softmax function:

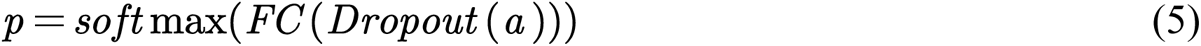

*FC* represents fully connected layer, the formula for a fully connected layer is:

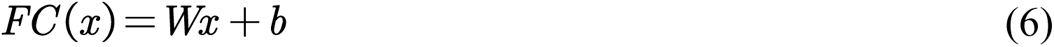

where *W* is the weight matrix, *r* is the input vector, and b is the bias vector. Dropout is a regularization technique used to prevent overfitting in neural networks. The formula is:

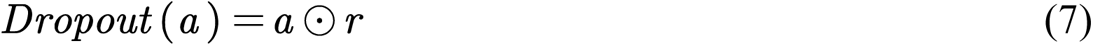

where *a* is the input vector, *r* is a vector of the same dimensions as *a*, with elements being either 0 or 1, sampled according to the retention probability *p*, and ⊙ denotes element-wise multiplication. The loss function is the cross-entropy loss function, defined as:

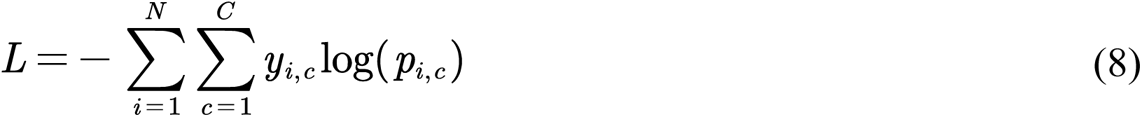

where *N* represents the total number of samples, *C* represents the number of subtypes, *y_i,c_* represents the indicator variable that sample *i* belongs to subtype *c* and *p_i,c_* is the probability that sample *i* is predicted to belong to subtype *c*.

The primary challenge we face is the imbalance in sample sizes across subtypes in the dataset, which can significantly impact classification performance, particularly for rare subtypes. To address this issue, we divide the subtypes into two groups — one with a large number of samples, including I-A, I-B, I-C, I-D, I-E, I-F, I-G, II-A, II-C, III-A, V-A, and the other with fewer samples, including I-U, II-B, IV-A3, V-B1, V-F1, V-F2, V-K, VI-A, VI-B1, VI-B2, VI-D. We employ a stacking strategy, training separate models for each group of subtypes, and using the output of each model as new features for the meta-model (XGBoost). Additionally, the original features (*k-mer* frequency, repeats length, GC content) are combined with these new features as inputs to the meta-classifier (XGBoost). However, during the training of the model for the second group of subtypes, the limited sample size impedes the model from adequately learning, resulting in suboptimal performance. Therefore, we employ a transfer learning strategy, fine-tuning pre-trained models, to transfer the knowledge learned from the pre-trained model of the first group of subtypes to the model training of the second group of subtypes. This strategy aims to enhance the model’s performance in scenarios with limited samples, leveraging the rich sample information from the first group of subtypes and transforming it into effective learning for the second group of subtypes, thereby improving the model’s generalization ability and classification performance.

### Performance evaluation

We evaluate CRISPRclassify-CNN-Att using weighted precision, recall, and f1-score. The precision measures the correct predictions among all instances predicted for a specific class, defined as follows:

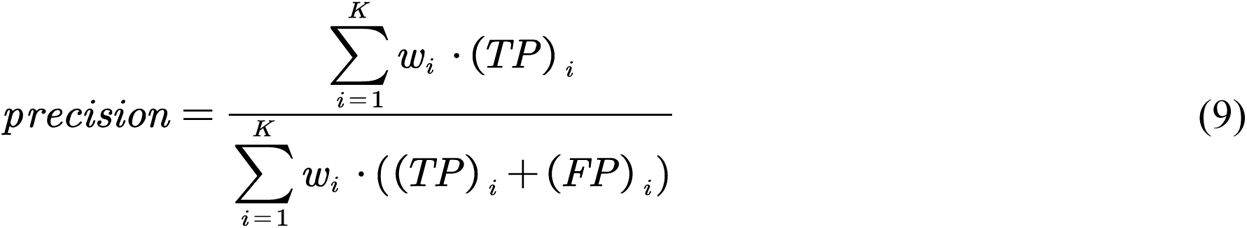

The recall measures the proportion of actual instances of a specific class that are correctly predicted, defined as follows:

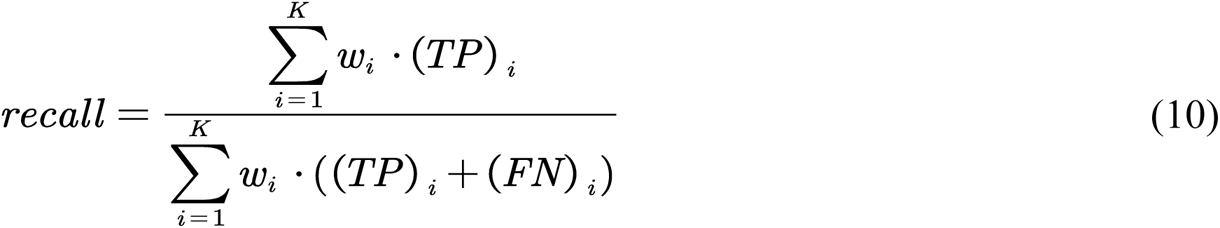

The f1-score is the harmonic mean of precision and recall, used to evaluate the overall performance of the model, defined as follows:

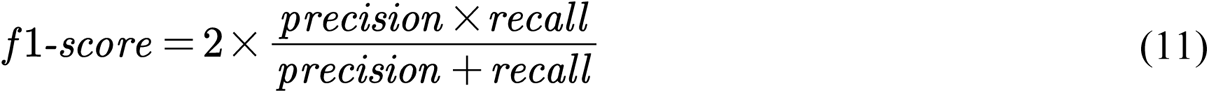

Where *K* is the number of classes, *w_i_* represents the weight of class *i*, (*TP*)_*i*_ and (*FP*)_*i*_ denote the number of true positives and false positives, respectively, for class *i*. (*FN*)_*i*_ denotes the number of false negatives for class *i*.

### Feature analysis

Feature analysis includes identifying important *k-mers* and analyzing the distribution of repeats length and GC content. We use the base model to analyze and identify important k-mers in each CRISPR-Cas system subtype. We systematically set the occurrence frequency of each *k-mer* in the test set to zero and observe changes in the model’s prediction metrics, including precision, recall, and f1-score for each subtype. A *k-mer* is considered important for a subtype if setting it to zero results in a decrease in prediction performance. Additionally, biological features such as sequence length and GC content significantly influence model performance. We calculate the distribution of these features for each subtype to assess their impact.

## Supporting information

CRISPRclassify...4-6-27-v5.pdf

