## Supplementary material for "Deep Learning-Based Classification of CRISPR Loci Using Repeat Sequences": CRISPRclassify...4-6-27-v5.pdf

**Table S1. Comparison results between CRISPRclassify-CNN-Att and CRISPRclassify methods**

| | **CRISPRclassify-CNN-Att** | | | **CRISPRclassify** | | |
|---|---|---|---|---|---|---|
| | **precision** | **recall** | **f1-score** | **precision** | **recall** | **f1-score** |
| **I-A** | 0.75 | 0.73 | 0.74 | 0.67 | 0.49 | 0.56 |
| **I-B** | 0.88 | 0.92 | 0.90 | 0.83 | 0.93 | 0.87 |
| **I-C** | 0.98 | 0.98 | 0.98 | 0.95 | 0.96 | 0.95 |
| **I-D** | 0.82 | 0.73 | 0.77 | 0.80 | 0.87 | 0.83 |
| **I-E** | 0.99 | 0.99 | 0.99 | 0.96 | 0.98 | 0.97 |
| **I-F** | 0.99 | 0.99 | 0.99 | 0.97 | 0.97 | 0.97 |
| **I-G** | 0.95 | 0.94 | 0.94 | 0.83 | 0.86 | 0.85 |
| **I-U** | 0.93 | 0.94 | 0.93 | 0.65 | 0.51 | 0.57 |
| **II-A** | 0.95 | 0.98 | 0.96 | 0.93 | 0.95 | 0.94 |
| **II-B** | 0.97 | 1.00 | 0.98 | 0.92 | 0.68 | 0.78 |
| **II-C** | 0.92 | 0.84 | 0.88 | 0.88 | 0.81 | 0.84 |
| **III-A** | 0.89 | 0.90 | 0.90 | 0.87 | 0.83 | 0.85 |
| **IV-A3** | 0.82 | 0.74 | 0.78 | 0.93 | 0.63 | 0.75 |
| **V-A** | 0.97 | 0.99 | 0.98 | 0.97 | 0.97 | 0.97 |
| **V-B1** | 1.00 | 0.56 | 0.72 | 0.70 | 0.93 | 0.80 |
| **V-F1** | 0.81 | 0.90 | 0.85 | 0.75 | 0.30 | 0.43 |
| **V-F2** | 0.85 | 0.85 | 0.85 | 0.96 | 0.73 | 0.84 |
| **V-K** | 0.97 | 0.91 | 0.94 | 0.85 | 0.82 | 0.83 |
| **VI-A** | 0.94 | 0.94 | 0.94 | 0.94 | 0.75 | 0.83 |
| **VI-B1** | 0.85 | 0.85 | 0.85 | 0.81 | 0.58 | 0.68 |
| **VI-B2** | 1.00 | 0.92 | 0.96 | 0.61 | 0.52 | 0.56 |
| **VI-D** | 1.00 | 0.94 | 0.97 | 1.00 | 0.80 | 0.88 |
| **All** | 0.940 | 0.941 | 0.940 | 0.911 | 0.912 | 0.911 |

**Table S2. The comparative results of feature ablation studies for Model-large and Model-less, using only repeat sequences and only extracted features**

| | Model-large | | | Model-less | | |
|---|---|---|---|---|---|---|
| | Model-large | Repeat | Extracted | Model-less | Repeat | Extracted |
| **precision** | 0.942 | 0.919 | 0.917 | 0.918 | 0.891 | 0.880 |
| **recall** | 0.940 | 0.919 | 0.919 | 0.920 | 0.899 | 0.883 |
| **f1-score** | 0.941 | 0.918 | 0.917 | 0.920 | 0.895 | 0.881 |

**Table S3. The important *k-mers* for each subtype based on encoding each *k-mer* in the sequence as [0, 0, 0, 0]**

| | | | | | | | | | | |
|---|---|---|---|---|---|---|---|---|---|---|
| **I-C** | AAAATC | AAAATG | AAACCT | AAATCA | AAATGG | AACCTT | AATCAA | AATGGA | ACATTG | ACTAGT |
| ACTTGG | AGCACG | AGTGAT | ATAATG | ATACTA | ATGGAG | CAAACC | CATTGA | CCACAC | CCCACA | CCTTAC |
| CTATAC | CTGAAA | CTGGCA | GACCCA | GCTGAC | GCTTCA | GGCTGA | GTGATA | GTGGAA | TAATGA | TCAAAA |
| TGAAAA | TGATAA | TTCAAA | | | | | | | | |
| **I-B** | CCACTC | TTCCAA | ACTTTC | CGAGGC | AAGCTC | AGAGAC | CATCCA | TCTCAA | CCATAA | GACCCA |
| TACAAA | GGTTGA | ATTTCC | TAGGAA | AATTGA | CAATCG | ACCCCT | AGAACT | CCCAAC | TAAGGA | ATTACG |
| ACGTAT | CCATCC | GAATTA | TATTAA | CCTATC | GTATAC | TTATAA | ATCTAG | TAACAA | ACAGCT | AGAGAT |
| GGCGGA | CATAGG | CCCTGA | CCCCTC | GTAGAC | ATGGAA | TAAAAA | CTACAG | ATTTAC | AACGCG | ACAGGT |
| CTCGTC | TGCACA | CTGGCA | AGGGAT | AGGCTA | AAAGAG | GTCTAA | GACATC | AATGAA | GATTGA | CGGTCC |
| GCCGAA | ACTACC | AGGGCG | GTCTCA | GATTCA | AGTCTC | ATTATC | AAACTT | ATGCCG | ATTAAT | AGTAGG |
| CAATAG | CCATAC | CCTACC | AGTTTC | ACGGGC | AGAACG | CTGACC | ATTAAA | ATTCCC | AAAACT | ATGCTC |
| ACAAAT | ATTCCA | GGGAAC | GAGGCA | CTCTGC | CTGAAG | GTAGAA | CTACTA | GCGGAC | AGTACC | CTGGGA |
| ATGAAC | GATTCC | GGCTAA | CAACGG | ACTTAG | CGCTAC | AACTTA | TTCAAA | AGACTC | ACGTGA | TTAAAA |
| CAAAAG | CTATTA | GGGGCA | ATTAGC | TGGAAA | CATCAC | AAAAGA | CCAGTA | AGTCCC | CGCTAA | ATGTTC |
| CTAAAG | CCTGAA | AAGATT | CTTATC | CACGCA | ATCCAG | CGATAC | AATTGT | CAGGCA | ATTAAC | CGCACC |
| **I-C** | CACGGA | GCGCCA | AAGGGC | CACCAC | ACGTGG | AATTAA | CCTGCC | GCCAGC | ACGAAC | GGATCA |
| ACGCGC | AGCCTC | CTGAGC | GACTGA | TCCGCA | GACGAA | GTGGCA | ACACAG | CGCCAC | CATCCA | ATTCTC |
| TCTCAA | GTAGCA | CATCAA | CGCACG | GGTTGA | GCACCA | GTCGCA | GCATCC | CCTCGC | CGCTAC | GCAACC |
| TTGAAA | AATTCT | AGGATC | GGACCA | GGGCGA | ATTAAG | CCGGCA | AGGGGC | CCGCCC | AGCGCT | CTGCTA |
| AGAAAT | AAGAGG | CCAAGC | CCCCGC | ATTGAA | CGAAGA | TTAAAA | CTACCA | AGAAAC | CCACGC | CCCCAC |
| CCGAGC | AGCGAT | AGAGGG | CGCTGC | CTCAAC | GGATGA | CAATTA | CCCACA | GAGTGC | CCCCTC | GCTGGA |
| ACTGAG | AGGGTG | ACCCCA | GAGTAA | ACAGGT | CCAGTA | GCATCA | GCACCC | CTCGTC | AATACC | AGCATC |
| GTCGAA | AAGCTT | CAATTC | CAGCGC | GGCACC | GATCGC | GTGCGA | ATCCAG | CACCCC | GCGCTA | AGCGAC |
| ACCAAG | ACCCAC | AGCGCC | ACCTGC | AACGTG | CGCACC | AGTAAG | GATCAC | CGGATC | GATTGA | AATTCC |
| CACGCG | ATGCGA | CATCCC | ATCCCC | GATTCA | CGAGCA | AGTCTC | CCCACG | GTCTCA | CACACA | CTAAAA |
| GTTGAA | CCAGCG | ATCCTC | CACAGG | GATTAA | GTAAGA | CGATCC | CGTGGA | ATCCGC | GCTACC | AGTGAT |
| ACGGGC | CAATCC | GAGGGC | ATTAAA | CGCGAC | ATCGCA | ATTCCA | GGAGAC | CCCTCG | ATCCTG | ATCACC |
| **I-D** | CACCCA | CTCCTA | TTCCAA | AAATCC | CAAAAC | CGAGGC | CTGAGC | GACTGA | CCACAC | AAGCTC |
| ACACAG | TAGGGA | GTTTCA | ATTATG | GCTCAA | CATGAG | AGCTCA | CCCTCC | TCTACA | ATTTCC | AATCCC |
| AATTGA | AAATTG | CAATCG | AACCCG | CAACGG | TTGAAA | GCTAAA | GGGTTA | CCCCGA | CCTATC | TAGCGA |
| AAGAGG | CCAAGC | ATTGAA | CTCTCC | CGAAGA | TCTGCA | TTAAAA | ACAACG | CAAAAG | AATCTA | ACCCAA |
| CAGTCC | AGAGAT | ACTAAA | CCCGGG | AGCGAT | AGAGGG | AAACCC | ATGCGG | GGATGA | CTTTCA | AAAAAT |
| CCCACA | CCCTCA | CCCCTC | ACTGAG | ATGAAA | TAAAAA | AAAAAC | CTGCAA | ACAGGT | AAAAG | GCACCC |
| CTGGCA | CGCTAA | ATGTTC | GAAACC | GTCGAA | AGGGAT | GGCACC | CATGCG | AACTAA | CCAAAA | AAAGA |
| AGCGAC | CCTCCC | ACCCAC | AATGAA | ACGGAA | AAACTC | CCTCAA | GATTCA | CACACA | ACTCTC | CTAAAA |
| GGATTA | CACAGG | CCCAAA | AAAATC | CAGGGA | AGTTTC | CAATCC | GAGGGC | ATTAAA | CGCGAC | AAATCT |
| AAAACT | CAAAAA | ATCTAC | CCCTTA | GGGAAC | GAGGCA | AAGAGA | CTACAA | CTCAAA | TCAAAA | AGGGA |
| **I-E** | GCGCCA | CGATTC | GGGGTA | ATGCGC | GCCAGC | ACGCGC | CGCGTA | AACCCC | CGCTTC | CACGGC |

| | | | | | | | | | | |
|---|---|---|---|---|---|---|---|---|---|---|
| AAGACA | CGCCAC | ACGGGA | CATGAG | ATTCTC | CCCTGC | TCCTCA | GAGCAC | GTCGCA | AATTGA | CAGCTC |
| CCTGCG | TTGAAA | ACCCCT | AGGAGC | AATTCT | GCAACC | CTACCC | AGGGGC | AGCGCT | AGAAAT | GGGTAC |
| ACCGCC | AGACAG | AGCAAC | ATTGAA | CCCCGC | GGAGCA | GCGGCC | CTCTCC | CGAAGA | CGCATA | ATGACG |
| CATTCA | GGGGGA | CCACGC | CCCCAC | AGATTG | ACTAAA | GGGGCA | CGCTGC | CAGGGG | AAGCGG | AACCGC |
| GAAGAC | CTCATA | CCTGCA | CAACCC | ACCCCG | CATCAC | CTTCCC | ACTCTG | GGGGAA | AATCCG | CCCGTA |
| AGGGGG | GACCCC | ACCCCA | AGCGGG | AGCTCG | GCACCC | CCACCC | CAATTC | CAGAGG | CAGCGC | CACGCA |
| CACCCC | GACTAC | GGGTCA | AAGACC | AGCGAC | CGGGGA | CCCAGC | ATTCAG | AGCGCC | CCGTAC | CTGCGC |
| CCCCCC | ACCTGC | CATATG | CGCACC | ACCCCC | GGTAAC | CACGCG | GGGTGA | CCCCCG | CATCCC | ATGCAG |
| CGGGAA | GATTCA | CCCACG | ACTCTC | ACGAGG | CCAGCG | ATCCTC | CGATCC | CGCGCG | AGTTCC | CCTACC |
| AGTGAT | ACTACA | AGGGTA | GTCCAC | GGGAAC | ACAGGA | CTACAA | CTCTGC | ACGCAG | GAAAAC | CCCGCA |
| ATCACC | | | | | | | | | | |
| **I-F** | GCAGTA | CAGCAG | ACATCA | GCCACC | GCCGAA | GGCACA | CCACCA | ATTATC | CGGCAC | ACGGCA |
| ACGTAT | GCAGCA | ACCGCC | GCACAC | ATGCCG | CAGTAG | CACGGC | GTATAC | AATGCC | ACAGGC | AAGCCA |
| AGCCAC | CCGTCA | CGTATA | CAGGCA | AGGCAG | CAATGC | ACCGTC | GACGTA | ATGAAC | | |
| **I-G** | AAGGGC | CACCAC | GAGTGC | ACGTGG | CCCCTA | CAGACA | GGATCA | ACGCGC | GACTGA | CGCGTA |
| AGTCCC | GTGGCA | CCTGAA | GGCACC | GATCGC | CTTATC | AAGCCA | AGCCAC | ACTCCA | CGATAC | CGGGGA |
| CCCCCC | TCACCA | TCCTCA | GGAAAC | AACGTG | CATATG | GATCAC | CCCGCA | CGGATC | CCTCGC | AGGATC |
| GCCACC | GCCTCC | GGGTGA | CCACCA | GGACCA | CTACCC | ACATGG | GACTCC | CCGCCC | GTTGAA | ATAGGG |
| CCCCGC | ATCCTC | GCGGCC | CGCATA | CGCGCG | CGTGGA | GGGGAC | CAGTCC | AAGCCT | ATCGCA | ATGAGG |
| CTCATA | AGCCTG | AATGAG | ACCCCG | CTTCCC | GCCTGA | ATCACC | | | | |
| **I-U** | AAAATG | AAATGG | AATGGA | ATGGAG | CCCTTC | CCTTCA | CGCCGC | CTGAAA | CTTCGA | TGAAAA |
| **II-A** | GAAGAA | CCTCAG | ACCAGC | AACACA | TTCCAA | AAACAA | GTGAGA | TAAATA | AATAAC | AATGGT |
| CATAAC | CAAAAC | AGAGGC | AACCCC | AAGCTC | AAGAAT | ACACAG | GTGTTA | ATTATG | TAATGA | GATTTA |
| CTTTAA | GACCCA | ATTCGA | GGAAAC | TTACAA | AAACTG | TAGGAA | AAATTG | GTTAAC | TTGAAA | ACCCCT |
| ATGTTA | TATTAA | GAATTA | AGAAAT | AAGAGG | ATCATA | CGACAC | ATACCC | ATCTAG | ATGACG | TCTAAA |
| ACAGCT | CCCCAA | GAATGA | AATACA | TGTAAA | AAAATA | AGAACC | CGAAAC | AGAATG | ATTGTC | AAATTA |
| CCCCTC | GTTTTA | AAAGCT | AACCTA | ACCCCA | CAAAGC | CAGACA | ATTTAC | CACCAA | CATGCA | TCAGAA |
| CGACCC | CAGAAA | ACTCAA | ACCTTC | AGTTAA | ATAAAA | AATTAG | CTCAGA | ACAACA | CCAAAA | CAGAAG |
| ACCTGC | GTCGTA | CCTAAG | GACATC | AAGTAC | AATGAA | TAATCA | AAGTTA | ATAATC | AGATAT | AATATC |
| ATGCAG | ACCAAT | ACCCTC | ATTTAA | GTAAAA | ATGCAT | ATTAAT | AAGAAA | AGAAGA | ATAACA | TCGACA |
| CTGACC | CTTATA | ATTAAA | AAATAA | AAAACT | GTTCCA | ATCAGG | AATCAT | CATTGG | GTAGAA | TATAGA |
| ATGACC | ATAACC | AATTAA | ATCTAA | AACAAC | ACATCT | CCCCTA | CATAGC | CACACC | CATACA | CTCGCC |
| ATTTGA | CAACAC | AGTACC | AATAGG | AATAAT | CAGCTC | GAGATA | ACGTGA | GATACA | TTAAAA | AGCATA |
| CTAGGA | CACTTA | AGAAAC | TCTGAA | CTATGC | CATCTA | CTGCAC | CCCTCA | AAATGA | GCAGAC | ACGACC |
| CGTGAG | CATATA | AACAGT | AATACC | AATGAC | ATATGA | CTAAAG | ATAAAT | CCTTCG | TCCAAA | CACAGC |
| ATACAC | ATTAAC | AGGAAA | ACACCA | CATTCG | CATCCC | AACTCA | AAATAC | GTTAAA | AGAGCT | AATTTA |
| ACAGAT | ATTTTG | GATAAC | AGATTC | ATGTAA | AGCTGC | ATCAAC | AAATTT | AATAAA | ACAAAC | ATGAGA |
| AACTGC | ATTTCA | | | | | | | | | |
| **II-B** | AAACCT | AACCTT | ACTAGT | AGTAAA | AGTGAT | ATACTA | CAAACC | CCTTAC | CTATAC | GCTGAC |
| GGCTGA | GTGATA | GTGGAA | | | | | | | | |
| **II-C** | TATAGA | CACCCA | ACCAGC | AACACA | AATTAA | ACTTCC | GGCAAC | AACAAC | TATTCA | TTCCAA |
| AAACAA | AATGGT | CAAAAC | CATAGC | AACCCC | ACACAG | ATTTGA | ATTATG | CAACAC | ATAGGC | GATTTA |
| GCTAGA | TTACAA | CTGGGA | AATAGG | AATTGA | AATCCC | GAAACA | AAATTG | CAGGTA | CAGCTC | GGCTAA |
| AGTGGA | ACCCCT | GAGATC | GCTAAA | GATGCA | GAGATA | ACTGCA | AGATAA | GACTCC | CTAACC | CAGTTC |
| CTAGAA | AGACTC | ATTGAA | GATACA | TTAAAA | AGCATA | CTAGGA | TAACAA | ACCCAA | ACAGCT | CAGTCC |
| AGAGAT | ACTAAA | TCTGAA | CTATGC | CTTTGA | TCTTAA | ACAGAG | TGTAAA | AAAATA | CGAAAC | CTGCAC |
| AAATGA | GTTTTA | GCTGGA | GTAGAC | AATTCA | TAAAAA | CAGCCA | CATATA | ATTTAC | GCAGCC | AGTCCC |
| ATGCTA | GCACCC | TGCACA | AATGAC | CATGCA | GGCACC | AGTTAA | GGAGAA | CCTATA | ATAAAA | AGGCTA |
| GACTAC | ACAACA | CCAAAA | GAGACA | AATCCT | TCCAAA | CCCAGC | CCCCCC | CACAGC | GACAGA | ATTAAC |
| AAGTAC | AAGTTA | ATTTTA | CACCAG | AGATAT | CTTAAG | ATGCAG | CACAAC | TACTAA | ATCCCC | ACCCTC |
| GTTAAA | AGAGCT | AATTTA | ACAGAT | CTAAAA | TCCCGA | ATTTTG | GATAAC | AAACTT | ACTTTG | CCAGCG |
| CGGTTA | ATGCAT | ATCCTC | AGATTC | ATTCAC | ATGTAA | AACTTC | AGCTGC | ATAACA | CAATAG | AAATTT |
| CCCAAA | CCTCTA | AAGTGG | ACGGCG | AGTTTC | ACTACA | AATAAA | CAATCC | CTCTTA | CTTATA | AAAACT |
| AAATAA | GTTCCA | ATCCCA | ACCAAA | GGAGAC | CTACAA | CGCAAC | AACATA | TCAAAA | AGACAG | |
| **III-A** | CCACTC | ACCAGC | CCTCAG | AACACA | TAAGTA | AAACAA | TAAATA | CAAAAC | AAGAAC | AACCCC |
| TCCGCA | AAGCTC | GTTTCA | GTGTTA | AGAGAC | ATTATG | ACTCCA | CCCTCC | GAACAC | TCTACA | CCCTGC |
| TCCTCA | GGAAAC | GAGCAC | TACAAA | AAACTG | TAGGAA | AATTGA | GAAACA | AAATTG | GAGACC | TACCTA |
| GTTAAC | GCATCC | CCTGCG | AACCCG | GCAACC | TTGAAA | CCATTA | AGAACT | GACGAC | ATTACG | TAAGGA |
| ATTAAG | CCCCGA | CCATCC | GAATTA | TATTAA | AGCAAC | ATCATA | ATTGAA | CCCCGC | ACACGC | AACCCT |
| ATGACG | GTAGGA | TCTAAA | CAGTCC | ACAGCT | AGAGAT | ACTAAA | CCCCAA | CCGAGC | AGTATC | GAATGA |
| GGCGGA | ACTGAA | ACAGAG | GGACGA | CATAGG | ATTCAA | AGAATG | GAAATC | CCCTGA | GTTTTA | CCCCTC |

| | | | | | | | | | | |
|---|---|---|---|---|---|---|---|---|---|---|
| CCCGTA | GTAGAC | AGGGGG | ATGGAA | AATTCA | AGGCAA | AGGGTG | ACCCCA | ATGAAA | AAAAAC | TAAAAA |
| AACGCG | GAGTAA | CACCAA | TCAGAA | CGACCC | CAGAAA | ATTTAG | CCTATA | GTGCGA | ATAAAA | CTCAGA |
| AGGGCA | GTCTAA | CACACG | CCTCCC | CTGCGC | ACCTGC | GACAGA | AATGAA | TAATCA | CTCCCC | ATAATC |
| GATTGA | GCTTGA | ATGCGA | ACTACC | AGCCCT | AGGGCG | CCCCCG | TACTAA | AAAGTC | ACCAAT | ACCCTC |
| ACGGTC | AATGGA | GATTAA | ATTCAC | AATATT | AGTAGG | CGCGCG | CCATAC | CCTACC | ATCCGC | AGAACG |
| AGTAAT | ATTAAA | ATTCCC | AAATAA | AAAACT | ACAAAT | GGAGAC | ATCCTG | CTGAAG | AATCAT | ACCCCC |
| AGACAG | ATAACC | GGCAAC | AACAAC | CCTGCC | GGGCAA | AGCCTC | ACGAAC | CTACTA | GACTGA | AAGTCT |
| CACACC | CATACA | GACGAA | GCGGAC | CATTAA | CGTAGA | CCCGAC | AATCCC | AATAAT | GATTCC | ACTTAG |
| AGGAGC | ACACAC | AACTTA | AGGAAT | GGGTTA | TTCAAA | GAGATA | AGATAA | AATTAC | AAAACA | GACTCC |
| GGAACA | CGTTGC | TAGCGA | TCGTCA | GGAGCA | ATATTA | TTAAAA | AGAAAC | GGGGGA | AGATTG | CCCGGG |
| GTTGCA | CTATGC | AAACCC | CGATGA | AGTCTA | CTCATA | TGGAAA | CATCAG | CAACCC | ACCCCG | CTGCAC |
| AAATCA | CCCTCA | AATCCG | ACGACC | CGCATC | AGTCCC | AATGAC | ATTGTG | GTTACA | GAAACC | CCACCC |
| **IV-A3** | ACAGGG | ATACAG | CAGGGA | CGTGTA | CTATAA | GTATAC | TATACA | | | |
| **V-A** | AGTTTC | ATAGAA | GCAGAC | GAGATC | AATAGA | CTATGC | AGTATC | CTACTA | CAGACA | ATTTTA |
| AGATCT | GCTTGA | | | | | | | | | |
| **V-B1** | AATCCG | AGATGA | ATCCGC | CAATCC | CCGCAC | CTGGCA | TCCGCA | | | |
| **V-F1** | ATGAAA | CTCCAC | AATCCG | GATGAA | CCGCAC | CAATCC | TCCGCA | AGATGA | ATCCGC | |
| **V-F2** | GATAAC | AGATAA | CTCCAC | ACAGGG | ATACAG | TATACA | GTATAC | CAGGGA | CGTGTA | CTGAAA |
| **V-K** | ACTTGG | TTGAAA | CCCACA | GTTGAA | GACCCA | CATTGA | TAATGA | ATAATG | CCACAC | ACATTG |
| TGATAA | | | | | | | | | | |
| **VI-A** | CCTTCA | GTGTAA | CAGTTC | AGGGAT | AGTTCC | CTTCGA | GTTTCA | CTATAA | CCCTTC | ATGGTC |
| **VI-B1** | AAAATC | AATCAA | AGCACG | TCAAAA | AAATCA | CTTCAA | TTCAAA | GCTTCA | | |
| **VI-D** | ATCACA | AATCAC | | | | | | | | |

**Table S4. Unique important *k-mers*, identified based on the negative impact on predictive performance when they are encoded as [0, 0, 0, 0] in the sequence, exist exclusively in individual subtypes**

| | | | | | | | | | | |
|---|---|---|---|---|---|---|---|---|---|---|
| **I-B** | ATACTC | AGGACT | AAGGAA | AAAATT | ATTAGC | GTTATA | CCCTAC | ACAGCC | CCATAA | CACTGA |
| ACTTTC | AATTAT | ACCCTA | ATGCTC | AGGCAT | CAGGAA | CCCAAC | ACAATA | GGCTTA | TTATAA | CTCGAC |
| CACCTA | CTATTA | GCTCAC | CGGTCC | CTACAG | | | | | | |
| **I-C** | GGGCGA | CGCACG | CACGGA | | | | | | | |
| **I-D** | ATGCGG | CTGCAA | TCTGCA | | | | | | | |
| **I-E** | AAGACA | GGGGAA | CATTCA | CGGGAA | AGCTCG | CGCTTC | ATTCAG | ACGGGA | AAGACC | GGGTAC |
| GACCCC | AGGGTA | GGGGTA | CCTGCA | ACAGGA | GAAGAC | AACCGC | GAAAAC | ATGCGC | | |
| **I-F** | CGGCAC | ACGGCA | ACATCA | GCACAC | GGCACA | | | | | |
| **I-G** | AGCCTG | GCCTGA | ATAGGG | AAGCCT | ATGAGG | GGGGAC | TCACCA | AATGAG | | |
| **I-U** | CGCCGC | | | | | | | | | |
| **II-A** | ATACCC | AGAACC | ACAAAC | CCTTCG | AACTCA | AACAGT | AATTAG | ACTCAA | ATACAC | CATTCG |
| CACTTA | ACCTTC | GTCGTA | GAAGAA | CTCGCC | ATTTCA | ATCAGG | AATACA | AAATAC | AATATC | AGAGGC |
| AAGAAA | CAGAAG | AACCTA | AATAAC | AGAAGA | CCTAAG | AAGAAT | | | | |
| **II-B** | AGTAAA | | | | | | | | | |
| **II-C** | CCTCTA | AATCCT | CTCTTA | CAGCCA | GATGCA | GGAGAA | AAGTGG | GCAGCC | TCTTAA | CGGTTA |
| CTTAAG | CTAACC | CACCAG | AGTGGA | ACGGCG | TATTCA | | | | | |
| **III-A** | ACGGTC | AACCCT | CACACG | GGACGA | AGGGCA | AAAACA | GACGAC | ACCTAT | AAAGTC | GGCAAA |
| ACACAC | ACACGC | CGCATC | GTTGCA | GGGCAA | ATTCAA | AAGTCT | AGGCAA | ACTGAA | | |
| **V-A** | ATAGAA | AATAGA | AGATCT | | | | | | | |
| **VI-A** | GTGTAA | ATGGTC | | | | | | | | |
| **VI-B1** | CTTCAA | | | | | | | | | |
| **VI-D** | ATCACA | AATCAC | | | | | | | | |
